# *In vitro* and *in vivo* development of the human intestinal niche at single cell resolution

**DOI:** 10.1101/2020.01.31.928788

**Authors:** Michael Czerwinski, Emily M. Holloway, Yu-Hwai Tsai, Angeline Wu, Qianhui Yu, Josh Wu, Katherine D. Walton, Caden Sweet, Charlie Childs, Ian Glass, Barbara Treutlein, J. Gray Camp, Jason R. Spence

## Abstract

The human intestinal stem cell (ISC) niche supports ISC self-renewal and epithelial function, yet little is known about the development of the human ISC niche. We used single-cell mRNA sequencing (scRNA-seq) to interrogate the human intestine across 7-21 weeks of gestation. Using these data coupled with marker validation *in situ*, molecular identities and spatial locations were assigned to several cell populations that comprise the epithelial niche, and the cellular origins of many niche factors were determined. The major source of WNT and RSPONDIN ligands were ACTA2+ cells of the muscularis mucosa*. EGF* was predominantly expressed in the villus epithelium and the EGF-family member *NEUREGULIN1* (*NRG1*) was expressed by subepithelial mesenchymal cells. Functional data from enteroid cultures showed that NRG1 improved cellular diversity, enhanced the stem cell gene signature, and increased enteroid forming efficiency, whereas EGF supported a secretory gene expression profile and stimulated rapid proliferation. This work highlights unappreciated complexities of intestinal EGF/ERBB signaling and identifies NRG1 as a stem cell niche factor.

## INTRODUCTION

The stem cell niche within a tissue is required to regulate stem cell maintenance, self-renewal and differentiation (Scadden, 2006). The niche is made up of both physical and chemical cues, including the extracellular matrix (ECM), cell-cell contacts, growth factors and other small molecules such as metabolites (Capeling et al., 2019; Cruz-Acuña et al., 2017; Gjorevski et al., 2016). Understanding the niche within various tissues has been central to understanding how tissues are maintain homeostasis, and for understanding how disease may occur (Van de Wetering et al., 2002). Further, establishing proper *in vitro* niche conditions has allowed the growth and expansion of gastrointestinal tissue-derived stem cells in culture (Dedhia et al., 2016; Kretzschmar and Clevers, 2016). For example, through understanding that WNT signaling is important for maintaining intestinal stem cell (ISC) homeostasis (Muncan et al., 2006; Pinto et al., 2003; Sansom et al., 2004), blockade of BMP signaling by NOGGIN (NOG) promotes ectopic crypt formation (Haramis et al., 2004), and EGF is a potent stimulator of proliferation (Goodlad et al., 1987; Ulshen et al., 1986), it was determined that WNTs, RSPONDINs (RSPOs), NOG and EGF can be utilized to expand and maintain ISCs in culture as 3-dimensional intestinal organoids (Ootani et al., 2009; Sato et al., 2009, 2011). This same information has been leveraged to expand and culture human pluripotent stem cell derived intestinal organoids *in vitro* (Finkbeiner et al., 2015; Spence et al., 2011; Wells and Spence, 2014).

Although a wealth of information about the signaling environment within the ISC niche has informed our understanding of ISC regulation, and that this information has been leveraged to grow epithelium-only intestinal organoids (herein referred to as enteroids) *in vitro*, it is also clear that current *in vitro* systems are still not optimized to most accurately reflect the *in vivo* environment. Efforts have been ongoing to improve the *in vitro* physical environment through biomimetic ECM (Capeling et al., 2019; Cruz-Acuña et al., 2017; Gjorevski et al., 2016), and by adjusting signaling cues to more accurately reflect the *in vivo* niche (Fujii et al., 2018). More recently, single cell technologies have started to reveal unprecedented amounts of information about the cellular heterogeneity of human tissue and the ISC niche during health and disease (Kinchen et al., 2018; Martin et al., 2019; Smillie et al., 2019), and will undoubtedly yield substantial information about cell types and niche cues that regulate ISCs in various contexts.

Here, we set out to better understand the cellular diversity and niche cues of the developing human intestine using scRNA-seq to describe the transcriptional signatures and using fluorescent *in situ* hybridization (FISH) and immunofluorescence (IF) to define the location of cells that make up the ISC niche. We sought to interrogate the cellular source of known stem cell niche factors. We determined that a major source of WNT and RSPO ligands are most highly expressed in the *ACTA2/TAGLN+* smooth muscle cells of the muscularis mucosae that reside just below the proliferative crypt domains. We further determined that *EGF* is most abundantly expressed, not in the mesenchyme, but by the enterocytes of the villus epithelium, several cell diameters away from the proliferative region of the crypt. We identified a subepithelial population of cells that lines the entire villus-crypt axis, marked by a *PDGFRA^HI^/DLL1^HI^/F3^HI^* expression profile, and found that these cells express the EGF-family ligand *NEUREGULIN1* (*NRG1*). Given that *NRG1* is expressed in mesenchymal cells adjacent to the proliferative crypts, we tested the effect of both EGF and NRG1 on fetal human duodenal enteroids and found that EGF potently stimulated proliferation, while NRG1 supports enhanced cell-type diversity. A combination of NRG1 plus low concentrations of EGF supported enhanced cell-type diversity, improved stem cell gene expression, and markedly improved enteroid forming efficiency. Together, these results suggest that NRG1 is an essential niche cue and can be used to more accurately mimic the human ISC *in vivo* niche in enteroid cultures.

## RESULTS

### Interrogating the developing human small intestine with single cell resolution

Given that little is known about mesenchymal cell heterogeneity within the fetal human intestine, we aimed to better understand the mesenchymal cell populations that make up the developing human ISC niche. To do this, we obtained samples of human fetal duodenum starting just after the onset of villus morphogenesis (gestational day 47; 47d) with samples interspersed up to the midpoint (132d) of typical full-term gestation (280d) (Figure S1A-B). Major physical changes occur throughout this developmental window with rapid growth in length and girth and morphologic hallmarks are noted including the formation of villi and crypt domains within the epithelium and increased organization and differentiation of smooth muscle layers within the mesenchyme (Chin et al., 2017) (Figure S1A).

In order to capture the full complement of cell types that contribute to the developing human intestine, we dissociated full thickness intestinal tissue from 8 duodenal specimens ranging between gestational day 47d-132 and used scRNA-seq to sequence 3,248 – 9,197 cells per specimen. 37,058 total cells were used in the analysis after passing computational quality filtering (Figure S1B). From these 37,058 cells, each time point was randomly downsampled to contain between 3,000-3,200 cells per timepoint to avoid analytical bias from any overrepresented stage. Following dimensional reduction and visualization with UMAP (Becht et al., 2019; Wolf et al., 2018), we used individual known cell type marker genes, or gene sets applied to a cell scoring system, in order to identify major cell classes including epithelial, mesenchymal, endothelial, enteric nervous and immune cells (Figure S1D-E, see Methods). By using both individual genes and gene sets, we identified clusters 0, 1, 2, 4 and 7 as non-immune or vascular mesenchymal cells, which were then computationally extracted for further analysis (Figure 1), thus excluding epithelial, neuronal, endothelial and immune cells from downstream analysis.

**Figure 1.**
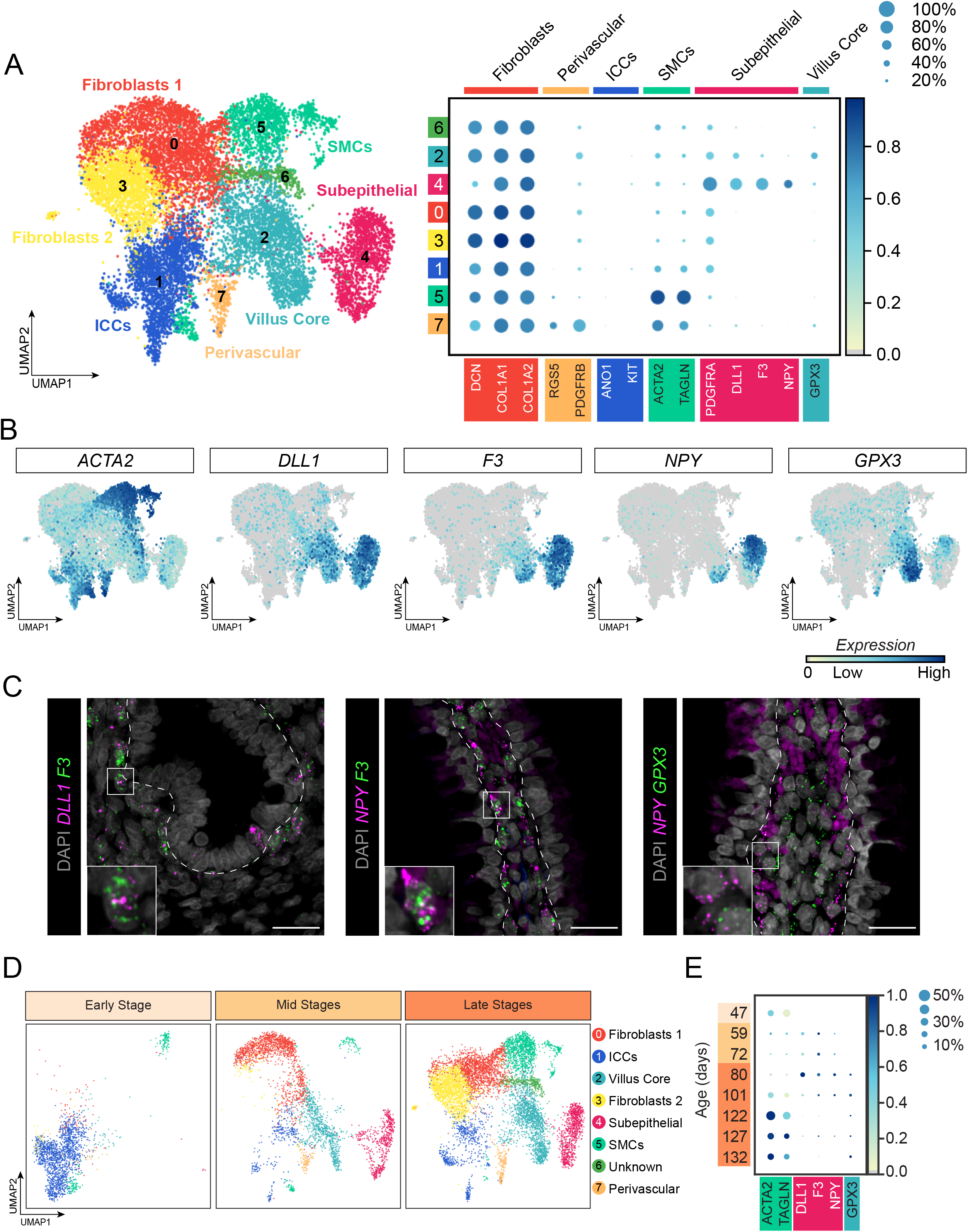
Mesenchymal heterogeneity in the developing human duodenum. **A)** UMAP plot of 13,847 computationally extracted mesenchymal cells identified 8 mesenchymal sub-populations, which were annotated using expression of known and unknown genes, shown in the dot plot (right panel). Dot size indicates the proportion of cells in each cluster expressing the gene, with the color indicating mean expression level (normalized z-score). **B)** Feature plots show that *ACTA2* and *DLL1* largely mark separate populations, and *DLL1* expressing cells can be further characterized based on expression of *F3, NPY* or *GPX3*. Cells with zero expression are shown in gray. **C)** Multiplexed fluorescent in situ hybridization staining of 132d human fetal intestine to show spatial localization of mesenchymal subpopulations (clusters 2 and 4). Left panel: *DLL1* (magenta), *F3* (green), DAPI (gray); Middle panel: *NPY* (magenta), *F3* (green), DAPI (gray); Right panel: *NPY* (magenta) and GPX3 (green) do not co-localize in villi, DAPI (gray). Dashed line defines the epithelial-mesenchymal boundary. Scalebars depicted represent 25μm. **D)** UMAP plots of cells separated by developmental time to assess the relative emergence of mesenchymal subpopulations, Early Stage (day 47) right, Mid Stages (days 59 and 72) middle and late stages (days 80, 101, 122, 127, 132) left. **E)** Dotplot of smooth muscle cells and villus mesenchyme markers during development. Dot size represents the proportion of cells in each cluster expressing the marker, with the color showing expression level (normalized z-score).

Re-clustering and UMAP visualization of extracted mesenchymal cells led to the identification of 8 predicted mesenchymal cell clusters (Clusters 0-7, Figure 1A). We began to assign cell identities to these clusters by examining genes known to be expressed in different intestinal mesenchymal cell populations, and by uncovering and validating new markers. We identified prominent clusters of cells (Cluster 0 and 3) that we identify as fibroblasts based on their expression of *COLLAGEN* genes (*COL1A1, COL1A2)* and *DECORIN* (*DCN*), combined with low-level expression of smooth muscle associated genes such as *ACAT2* and *TAGLN* (Figure 1A) (Kinchen et al., 2018). Perivascular cells (Cluster 7) were identified by expression of *RGS5* and *PDGFRB* as recently described via scRNA-seq analysis in the adult human intestine (Kinchen et al., 2018); Interstitial Cells of Cajal (ICCs, Cluster 1) were identified by *ANO1* and *KIT* expression (Gomez-Pinilla et al., 2009; Hwang et al., 2009); Smooth muscle cells (SMCs, Cluster 5) were identified by high expression of *ACTA2* and *TAGLN*; and we identified two population of cells that are defined broadly by the expression of the NOTCH ligand *DELTA-LIKE 1* (*DLL1*) (Clusters 2 and 4), which also expressed *PDGFRA* and *F3* (Figure 1A-B). *F3* was recently shown to be expressed in a population of mesenchymal cells that is adjacent to the human colonic epithelium (Kinchen et al., 2018). Clusters 2 and 4 were each enriched for unique markers as well, such as *NPY* (Cluster 4) and *GPX3* (Cluster 2) (Figure 1A-B).

In order to understand how cell populations identified in Cluster 2 (*GPX3+/PDGFRA^LO^/DLL1^LO^/F3^LO^*) and Cluster 4 (*NPY+/PDGFRA^HI^/DLL1^HI^/F3^HI^*) are spatially organized within the tissue, we used a combinatorial staining approach, utilizing multiplexed fluorescent *in situ* hybridization (FISH) and immunofluorescence (IF) to demonstrate that *DLL1*^*HI*^/*F3*^*HI*^ cells sit adjacent to both the crypt and villus epithelium (Figure 1C, Figure S2), with *NPY+* cell restricted to the subepithelial cells lining the villus (Figure 1C), but not that of the crypt (Figure S2). *GPX3*, which was expressed in the Cluster 2 *PDGFRA^LO^/DLL1^LO^/F3^LO^* cells, was most abundant in cells within the core of intestinal villi (Figure 1C). Cells expressing *GPX3* are observed sitting adjacent to *NPY*+ cells, and although scRNA-seq suggested that *GPX3+* cells express low levels of *F3*, *F3* mRNA was not abundant in the villus core via FISH (Figure 1C).

### Mesenchymal cell lineages emerge across developmental time

In order to get an idea of how gene expression patterns and cell populations may change over developmental time, we examined how different developmental time points contributed to each mesenchymal cluster (Figure 1D). We observed that cellular complexity appears to emerge and expand over time. At 47d, the *NPY+/PDGFRA^HI^/DLL1^HI^/F3^HI^* cells are entirely absent from the intestinal mesenchyme (Figure 1D, Cluster 4 - pink), and other cell populations (Clusters 0,2,3,5,6,7) make up a very small proportion of cells at this time, with the majority of 47d cells belonging to Cluster 1. Expression of *DLL1, F3, NPY* and *GPX3* were non-detectable at 47d (Figure 1E). *NPY+/PDGFRA^HI^/DLL1^HI^/F3^HI^* Cluster 4 cells first become apparent at 59d (Figure 1E), and robust *GPX3* expression is not observed until 80d or later. Based on the emergence of cell populations after 47d, these data suggest that mesenchymal heterogeneity increases over time, with cells after 80d distributing to all clusters (Figure 1D).

### Identifying putative human ISC niche factors in the developing gut

It has been demonstrated that several niche factors allow adult and developing human and murine intestinal epithelium to be cultured *ex vivo* as organoids (Capeling et al., 2019; Finkbeiner et al., 2015; Fordham et al., 2013; Hill et al., 2017; Kraiczy et al., 2017; Sato et al., 2009, 2011). These factors often include WNT and RSPO ligands, BMP/TGFβ antagonists and EGF, and are based on defined growth conditions that allow expansion of intestinal epithelium *in vitro* (Sato et al., 2009, 2011). Efforts have been made to determine more physiological niche factors for *in vitro* culture systems based on observed *in vivo* niche cues (Fujii et al., 2018), however these attempts at characterization have yet to fully leverage new high resolution technologies such as scRNA-seq. To identify putative niche factors we first determined which cells within the human fetal intestine expressed various known niche factors. We then identified the cellular origin of each niche factor *in silico* with scRNA-seq data, and validated these findings spatially in tissue sections using FISH/IF (Figure 2). We observed that *NPY+/PDGFRA^HI^/DLL1^HI^/F3^HI^* Cluster 4 subepithelial cells, and *GPX3+/PDGFRA^LO^/DLL1^LO^/F3^LO^* Cluster 2 cells that localize to the villus core lack robust expression of most known niche factors (Figure 2A). *RSPO2*, *RSPO3* and *WNT2B* are expressed most highly by the *ACTA2^HI^/TAGLN^HI^* Cluster 5 cells, and not expressed by the *PDGFRA^HI^/DLL1^HI^/F3^HI^* subepithelial Cluster 4 cells (Figure 2A, Figure S3), and expression of additional *WNT* and *RSPO* ligands was not detected. IF for the protein product of *TAGLN* (SM22), combined with FISH for *RSPO2*, *RSPO3* or *WNT2B* revealed expression of these ligands within the SM22+ cells located below the proliferative crypt/intervillus domain as well as by other fibroblast populations further away from the epithelium, consistent with scRNA-seq data (Figure 2B, Figure S3).

**Figure 2.**
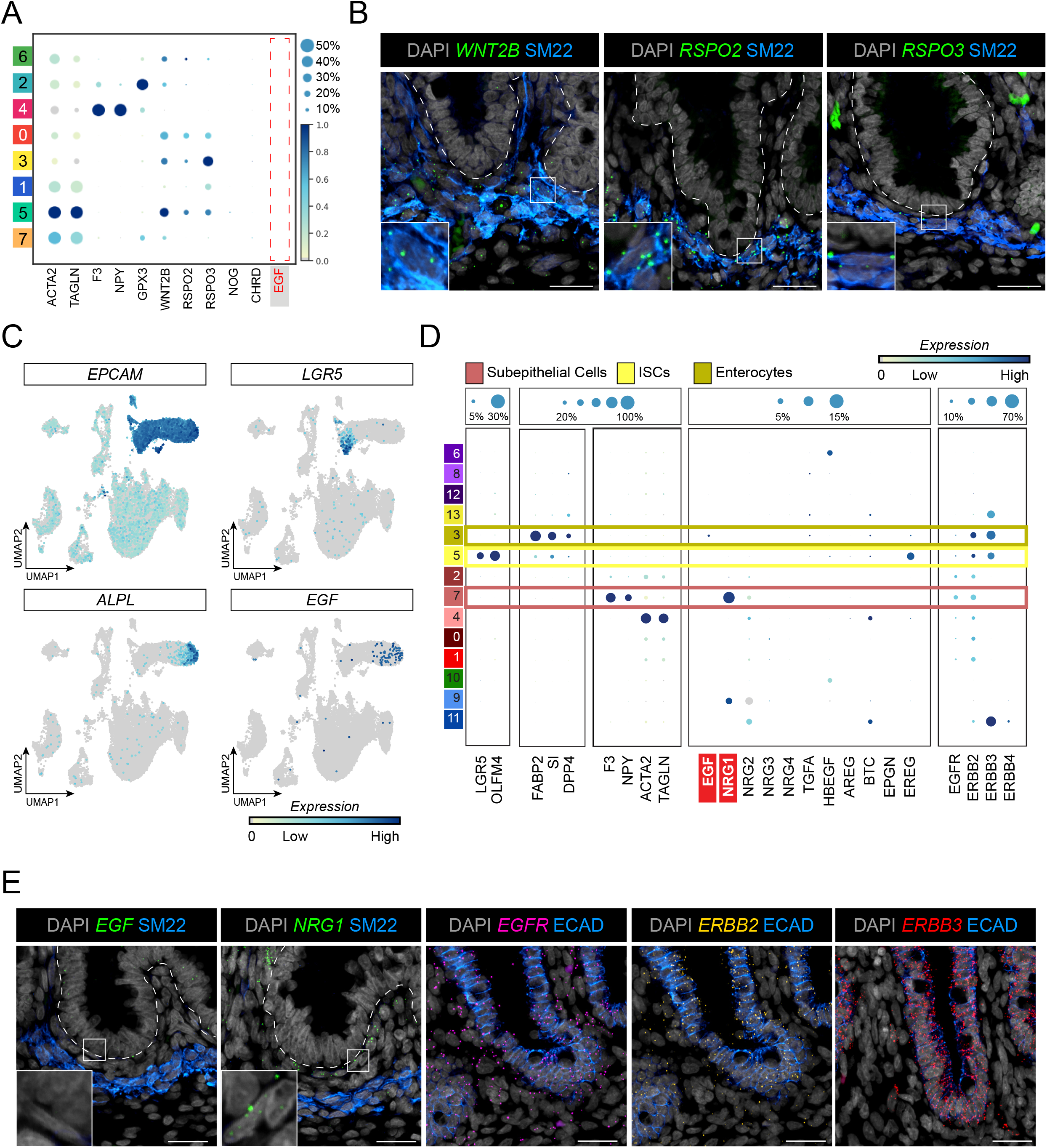
Interrogating stem cell niche factors in human intestinal mesenchyme. Dot plot showing smooth muscle, and villus mesenchyme markers alongside known ISC niche factors. Dot size represents the proportion of cells in each cluster expressing the marker, with the color showing mean expression (normalized z-score). Red box highlights the absence of EGF expression **B)** Co-fluorescent in situ hybridization and immunofluorescent staining of 132d fetal human intestine. *WNT2B*, *RSPO2* and *RSPO3* colocalize with SM22 protein (gene name *TAGLN*), a smooth muscle marker. Scalebars represent 25μm. **C)** UMAP plots of the entire data set (epithelium, mesenchyme, immune, neuronal, endothelial). Feature plots depicting the epithelial-specific (*EPCAM+*) expression of *EGF*. Within the epithelium, EGF is expressed by enterocytes (*ALPL*+) but is excluded from stem cells (*LGR5+*). Cells with zero expression are shown in gray. **D)** Dot plots showing gene expression of markers for stem cells (*LRG5*, *OLFM4*), enterocytes (*FABP2*, *SI* and *DPP4*), subepithelial mesenchyme (*F3*, *NPY*) and smooth muscle (*ACTA2*, and *TAGLN*) are shown alongside EGF family ligands and ERBB receptors. **E)** Co-fluorescent in situ hybridization and protein staining of 140d human intestine to determine the localization of EGF-family ligands (*EGF* and *NRG1*) and ERBB receptor expression. Protein localization of SM22 (blue) is shown alongside *EGF* and *NRG1* (green, left) and DAPI (gray). Dashed line defines the epithelial-mesenchymal boundary. Protein staining for ECAD (blue) marks the epithelium in FISH staining for ERBB family receptors *EGFR* (magenta), *ERBB2* (yellow) and *ERBB3* (red) and DAPI (gray) in each, right. All scalebars depicted represent 25μm.

The TGFβ-family inhibitors NOG and CHRD appeared at low levels with NOG slightly enriched β in *ACTA2+/TAGLN*+ smooth muscle cells by scRNA-seq. However, scRNA-seq data demonstrated more prominent expression of *GREM2*, and to a lesser extent *GREM1*, by the *ACTA2+/TAGLN+* population implicating them as the dominant TGFβ-family inhibitors β expressed in the human fetal intestine (Figure S3C). We also noted that *EGF*, was not robustly expressed within the mesenchymal population (Figure 2A). EGF is critical for epithelial proliferation *in vitro*, as inhibition of EGF has been demonstrated to induce a state of quiescence in LGR5+ ISCs in murine organoids (Basak et al., 2017). To further interrogate whether *EGF* is expressed in the developing intestine, or if other EGF-family members may be present, we examined expression of *EGF* and EGF-family members in the entire data set, including epithelium, immune, vasculature and enteric neurons (Figure S1B-D). We observed that *EGF* is expressed in a small subset of differentiated *EPCAM+/ALPL+* epithelial enterocytes (Figure 2C-D), a finding that was supported using co-FISH/IF and showed *EGF* is robustly expressed only in the villus epithelium (Figure 2E, Figure S2B), several cell diameters above the MK67+ crypt region (Figure S2D).

### ERBB pathway members are expressed in sub-epithelial niche cells

To investigate whether other EGF/ERBB family members are expressed, we surveyed expression of all known EGF-family member ligands and ERBB receptors in the single cell data (Figure 2D). We found that another ERBB ligand, *NRG1* is enriched in the *PDGFRA^HI^/DLL1^HI^/F3^HI^* subepithelial cells in both the villus and the crypt (Figure 2D-E). Co-FISH/IF supported the single cell data, showing that *EG*F is nearly undetectable in the crypt region, but that *NRG1* is expressed in sub-epithelial cells adjacent to the crypt (Figure 2E, Figure S2D). The ERBB receptors, including *EGFR, ERBB2* and *ERBB3* were expressed throughout the epithelium (Figure 2E).

### NRG1 enhances stem cell gene expression, and reduces proliferation associated gene expression

Based on the expression pattern of *NRG1*, we hypothesized that it may act as an ERBB niche signaling cue and may be physiologically relevant *in vitro* based on its localization and proximity to ISCs within the developing intestine *in vivo*. To interrogate the effects of NRG1 and EGF on the intestinal epithelium, we split human fetal duodenum derived epithelium-only intestinal enteroids in culture using standard growth conditions (WNT3A/RSPO3/NOG plus EGF, see Methods) into two groups. One group of enteroids was cultured in standard media with EGF (100ng/mL), the other was grown without EGF and was instead supplemented with NRG1 (100ng/mL)(Figure 3A). Following growth for 5 days in EGF or NRG1, enteroids did not appear phenotypically different (Figure 3A), but each group was subjected to scRNA-seq to investigate transcriptional responses between groups. Despite varying only EGF or NRG1 in the culture, we observed a strong shift in gene expression between the two groups as visualized in UMAP plots illustrated by near complete independent clustering of cells by culture media composition (Figure 3B). At the transcriptional level, NRG1 cultured enteroids increased stem cell marker gene expression including *OLFM4* and *LGR5* (Figure 3C-D), while proliferation associated gene expression (*MKI67, TOP2A*) was lower relative to enteroids grown in EGF (Figure 3C-D).

**Figure 3.**
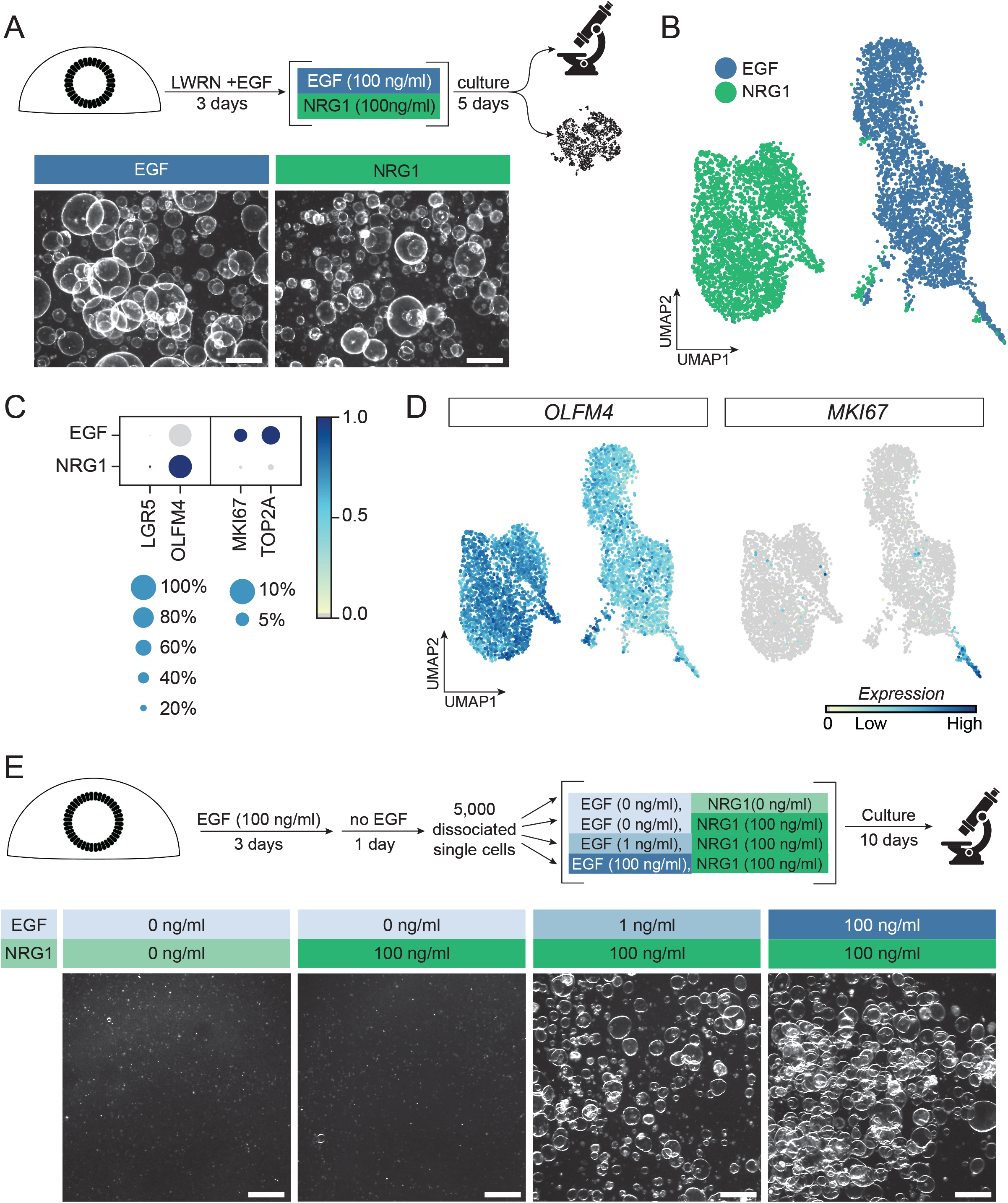
EGF and NRG1 drive strong transcriptional shifts in short term enteroid cultures. **A)** Schematic of experimental design (top). Stereoscope images of enteroids after 5 days of 100ng/ml EGF (left) or 100ng/ml NRG1 (right). B) UMAP plot of combined EGF-treated and NRG1-treated enteroids reveals treatment-dependent clustering. **C)** Dot plot depicting expression of stem (*LRG5, OLFM4*) and proliferation (*MKI67, TOP2A*) markers in EGF and NRG1 treated enteroids. Dot size represents the proportion of cells in each cluster expressing the marker, with the color showing mean expression level (normalized z-score). **D)** Feature plots demonstrating *OLFM4* and *MKI67* expression **E)** Schematic of experimental design (top). Stereoscope images of enteroids after single-cell passaging and growth with varying concentrations of EGF and/or NRG1. All scalebars depicted represent 500μm.

To functionally evaluate the observation that proliferation was reduced in NRG1 treated enteroids, we bulk passaged enteroids by fragmentation with a 30-gauge needle (see Methods) and then allowed them to expand for 3 days in standard (EGF 100ng/mL) growth conditions. We then removed EGF for 24 hours, dissociated enteroids into a single-cell suspension and plated 5,000 cells per droplet of Matrigel. Immediately upon seeding single cells, we added standard growth media supplemented with no-EGF (control), with EGF (100ng/mL) only, with NRG1 (100ng/mL) only or with NRG1 (100ng/mL) and a low concentration (1ng/mL) of EGF. After allowing single cells to expand for 10 days, we observed almost no enteroid recovery in the no-EGF control nor in the NRG1-only supplemented cultures. In cultures supplemented with EGF (100ng/mL) we observed robust growth, and adding just 1ng/mL EGF rescued the recovery defect seen in the NRG1-only group (Figure 3E).

### Long-term enteroid growth in NRG1 is associated with increased epithelial diversity *in vitro*

The previous experiment was conducted with enteroids that had been established and expanded in long-term culture with EGF, and the experimental data (Figure 3) suggested that these cultures were highly dependent on EGF. To determine the effects of different EGF-family members on establishment and long-term growth of enteroids, we cultured freshly isolated epithelium with EGF, NRG1 or a combination of EGF and NRG1 (Figure 4A). We used these cultures to carry out imaging, quantitative enteroid forming assays and scRNA-seq (Figure 4A). Isolated intestinal crypts were placed in Matrigel with culture medium containing no EGF/NRG1 (control), or containing NRG1 (100 ng/ml) plus increments of EGF (1-100ng/ml). Enteroids were successfully established from intestinal crypts under all conditions (Figure 4B). We noted that enteroids grown in high levels of EGF were phenotypically distinct from those grown in high NRG1 and low EGF, where high doses of EGF resulted in a cystic morphology, while NRG1 with zero or low (1ng/mL) EGF condition had much smaller and condensed morphology (Figure 4B-C). All conditions successfully underwent serial passaging, with the exception of the controls (no EGF/NRG1), which failed to expand beyond initial plating (Passage 0; P0) (Figure 4C). To determine the effects of different growth conditions on enteroid forming ability, we performed a quantitative single cell passaging assay on surviving cultures at P2. To do this, we dissociated the four remaining treatment groups into single cells and plated either 10,000 single cells (Figure 4D) or 1,000 single cells (Figure 4D) per droplet of Matrigel, allowed cultures to grow for 11 days, and quantified the number of recovered enteroids per 1,000 single cells (Figure 4D’). All groups included 100ng/mL NRG1, and groups with 0 and 10ng/mL of EGF led to a ~1% enteroid forming efficiency, whereas the 100ng/mL EGF group had ~0.5% enteroid forming efficiency and the 1ng/mL EGF condition led to a ~5.6% enteroid forming efficiency (Figure 4D’).

**Figure 4.**
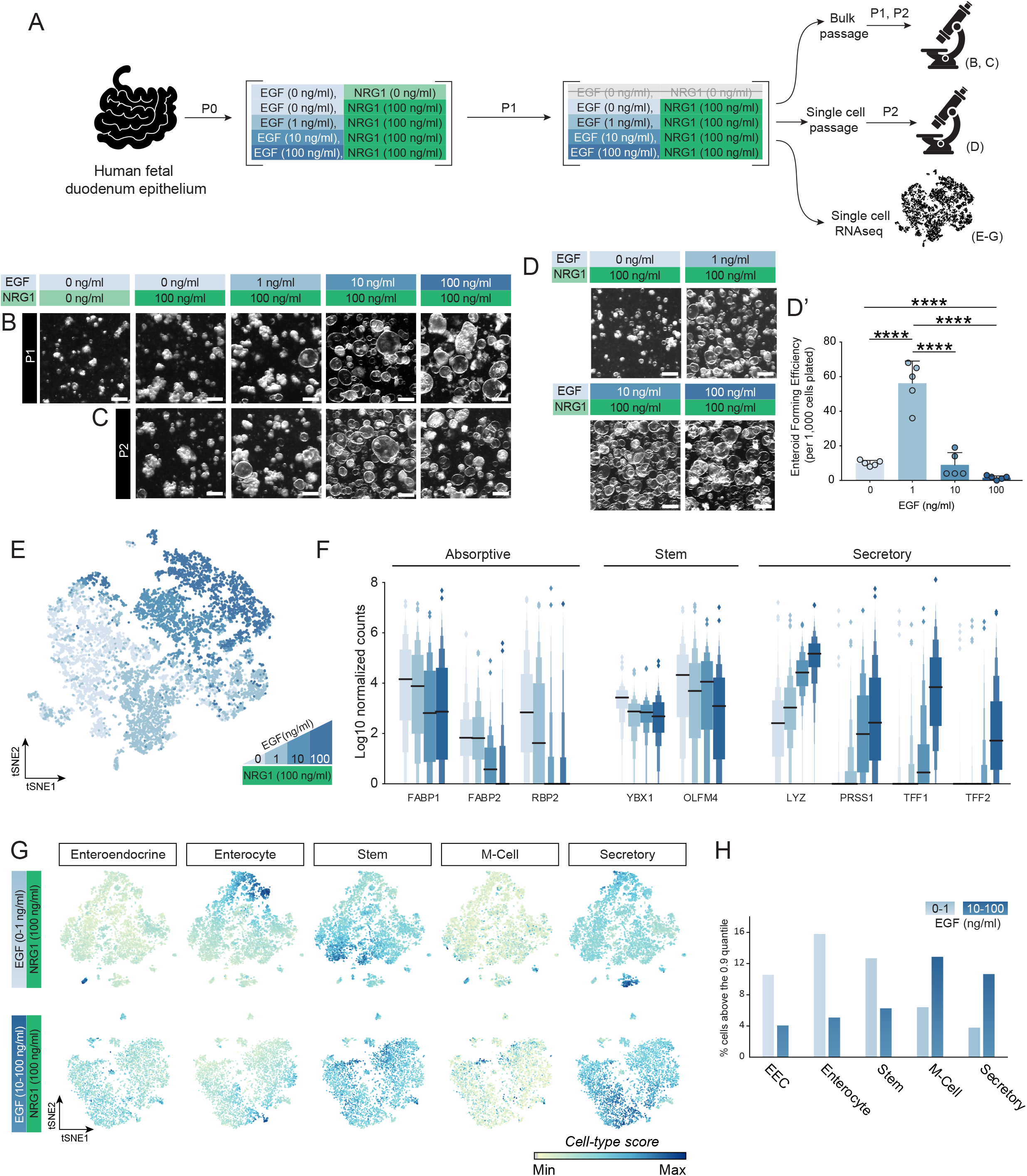
Establishment of new enteroid lines in NRG1 and low EGF increases cell type diversity *in vitro*. **A)** Schematic of experimental design. **B-C)** Stereoscope images of human fetal intestinal enteroid cultures established and grown in various concentrations of EGF with NRG1 after the first (P1) or second passage (P2). **D-D’)** Stereoscope images of 10,000 cells (D) and quantification of 1,000 cells(D’) of enteroid forming efficiency of P2 enteroids after single cell passaging in various concentrations of EGF. Data is plotted as the mean +/− SEM. Statistically significant variation in means was calculated using a one-way ANOVA (alpha=0.05) followed by Tukey’s multiple comparisons test of the mean of each group to the mean of every other group. Data are from a single experiment. Estimated p values are reported as follows: * p<0.05; ** p<0.01, *** p<0.001, **** p<0.0001. **E)** tSNE plot of scRNAseq results from enteroids grown in the presence of constant NRG1 (100 ng/ml) and varying concentrations of EGF (0, 1, 10, or 100ng/ml). **F)** Letter-value (Seaborn boxen) plot showing extended quantile data of gene expression for absorptive (*FABP1*, *FABP2*, *RBP2*), stem (*YBX1*, *OLFM4*) and secretory (*LYZ*, *PRSS1*, *TFF1*, *TFF2*) enriched genes across EGF treatment groups in the presence of NRG1 (100 ng/ml). **G)** tSNE plots depicting cellular diversity using cell-type scores for enteroendocrine, enterocyte, stem, M-cell, and secretory cells present in enteroids grown in low EGF (0-1ng/ml) and high EGF (10-100ng/ml) conditions. **H)** Percent of cells, per treatment group, that score at or above the 0.9 quantile of all cells in low EGF and high EGF treatment groups. All scalebars depicted represent 500μm.

Given the morphological differences between cultures grown in NRG1 with no/low EGF and high EGF, we wanted to interrogate the molecular differences between the groups. We therefore generated scRNA-seq data for each group, and sequenced 3,448 cells grown in 0ng/mL EGF, 3,405 cells grown in 1ng/mL EGF, 1,932 cells grown in 10ng/mL EGF and 1,884 cells grown in 100ng/mL EGF. tSNE dimensional reduction suggested that enteroids grown in low EGF (0, 1 ng/mL) clustered together, whereas enteroids grown in higher EGF (10, 100ng/mL) clustered together (Figure 4E). We further examined individual genes expressed in the various samples that are associated with the absorptive (*FABP1, FAPB2, RBP2*), secretory (*LYZ, PRSS1, TFF1, TFF2*) and stem cell populations (*YBX1, OLFM4*) as depicted in boxen plots (letter-value plots) (Hofmann et al., 2017) (Figure 4F). We observed that low-EGF conditions had higher expression of individual absorptive and stem cell markers whereas high-EGF conditions had higher expression of secretory markers, including those associated with the gastric epithelium (*TFF1, TFF2*) (Lennerz et al., 2010; Leung et al., 2002; Newton et al., 2000) (Figure 4F). By defining gene sets based on well-established ISC, absorptive enterocyte, secretory cell, enteroendocrine cell and M-cell marker genes (Haber et al., 2017), we combined low EGF (0-1ng/mL) and high EGF (10-100ng/mL) treatment groups and generated a score for each cell-type within each set of EGF treatment groups (Figure 4G). The high EGF group appeared to score homogenously for stem cell, absorptive and secretory genes across most cell clusters, whereas the low-EGF group appeared to contain more clearly distinct populations of enterocytes, enteroendocrine, stem and secretory cells (Figure 4G). In order to quantitatively determine if the low EGF group had a higher cell-type diversity, we examined the percent of cells from each group (low EGF vs. high EGF) in the 90^th^ quantile of each cell-type score (Figure 4H). These data supported the notion that enteroids cultured under conditions with NRG1 (100ng/mL) and low EGF (0-1ng/mL) possessed a higher proportion of enteroendocrine cells (EEC), enterocytes and stem cells, but had fewer secretory cells compared to enteroids grown in high EGF conditions.

Taken together, our data demonstrates that EGF strongly promotes proliferation in enteroids generated from the developing human intestine, but that at high doses (10-100ng/mL), induces reduced cellular diversity, with the majority of cells tending to skew towards the transcriptional signature of secretory cells, including genes that are normally associated with gastric secretory cells. In contrast, enteroids grown in NRG1 with low EGF have enhanced cellular diversity.

## DISCUSSION

### Subepithelial niche cells in the developing human intestine

We observed that *WNT2B*, *RSPO2* and *RSPO3*are expressed in the muscularis mucosae in the human fetal intestine, a muscle layer that is located close to the base of the epithelium. This is a unique finding when compared to recent single cell studies in the adult human colon, and compared to findings in the mouse. In the adult human colon, a source of WNT and RSPO external to the muscular mucosae has been identified as *WNT2B/RSPO3+* fibroblasts (Smillie et al., 2019), whereas in mice the predominant source of RSPO3 are PDGRFA+ cells in the small intestine (Greicius et al., 2018), or MYH11+ myofibroblasts in the colon (Harnack et al., 2019). In addition, the identification of the *Foxl1*+ telocyte has represented a major advance in elucidating the cells and sources of many niche factors in the murine intestinal stem cell niche (Aoki et al., 2016; Shoshkes-Carmel et al., 2018). When *Foxl1+* cells are genetically ablated, the crypt collapses, and these cells have been shown to be a major source of several niche factors, including WNT2B and WNT5A, and while they also express RSPO3, it was also demonstrated that non-telocytes also express RSPO3 (Shoshkes-Carmel et al., 2018). It remains to be seen if there is a unique expression pattern in the adult human small intestine and/or if there are changes in the cellular sources for *WNT2B* and *RSPO2/3* as development progresses; however, as the gross anatomical structure of the intestine observed in the fetal stages starting at 80 days of gestation and onward is maintained into adulthood (i.e. crypt-villus axis, muscle layers) it is possible that there are dramatic differences across species and regions of the gut for the major niche cells.

In the current work, we identify a subepithelial cell that lines the entire crypt-villus axis, marked by high levels of *DLL1*, *F3* and *PDGFRA expression*. These cells can be further sub-divided using the marker *NPY*, which is expressed in the villus (*NPY/DLL1/F3)* but not the crypt. Within the *DLL1/F3* transcriptional signature, we also observed robust expression of *FRZB* and *SOX6* (Figure S3), which have previously been described in the human colon as a subepithelial cell population that expresses several WNT family members (*WNT5A, WNT5B*) and several BMP family members (*BMP2, BMP5*) (Kinchen et al., 2018). Thus, while the focus of the current manuscript is on EGF-family members, it is likely that the niche signaling role for the *DLL1^HI^/F3^HI^* is more complex, and may involve secretion of activators and inhibitors of several other signaling pathways.

### Establishing signaling gradients along the crypt-villus axis

While difficult or impossible to test in human tissue, one could speculate that the robust levels of *EGF* expressed in the villus epithelium coupled with *NRG1* expression in subepithelial cells along the crypt-villus axis help to establish, in effect, a gradient of EGF/NRG1 signaling by differential receptor binding/dimerization in different domains. For example, high *NRG1* is expressed in the crypt-associated subepithelial cells, with low/no *EGF* being expressed in the in the crypt domain, whereas both NRG1 and lower levels of EGF are present in the putative TA zone, Finally, high NRG1 and high EGF is likely present in the villus, based on FISH data, but this area would also have the lowest levels of RSPO and WNT, given our localization data showing that ACTA2+/SM22+ cells of the lamina propria express undetectable levels of these genes, and that these ligands are produced in ACTA2+/SM22+ cells near the base of the crypt. In an attempt to mirror these *in vivo* expression patterns, *in vitro* experiments suggest that varying levels of NRG1 and EGF can drive different enteroid phenotypes, cellular diversity and stem cell function (enteroid forming efficiency) in conditions replete with WNT/RSPO/NOG. Given that *EGF* expression is highest in the villus epithelium, one might speculate that EGF normally acts as a differentiation factor given its expression *in vivo* coupled with our data showing that enteroids grown in >10ng/mL EGF had a molecular profile that was shifted toward a secretory lineage, including expression of Trefoil factor (*TFF)* genes canonically associated with the gastric epithelium.

Taken together, our data reveals that the human fetal intestinal stem cell niche is composed of multiple cellular sources, and highlights a unique role for different ligands from the EGF family. The resources we provide here lay the groundwork to further interrogate cellular relationships in the human fetal intestine, provide an important benchmark for *in vitro* experiments, and will inform additional methods to generate more robust and physiologic culture conditions.

## Supporting information

Supplemental File - Cell Scoring Gene Lists

## Acknowledgements

We thank Judy Opp and the University of Michigan Advanced Genomics Core for their expertise operating the 10X Chromium single cell capture platform and sequencing expertise. We would also like to thank the University of Michigan Microscopy Core for providing access to confocal microscopes and image analysis software, and The University of Washington Laboratory of Developmental Biology staff.

## Author contributions

MC and JRS conceived the study. JRS supervised the research. AW, EMH, YHT, MC developed tissue dissociation methods and generated single cell RNA sequencing data. MC, JW, QY, BT and GC performed computational analysis. MC, EMH, JW, QY, BT, GC and JRS interpreted computational results. EMH, KDW, CS, CC performed FISH experiments and imaging. YHT and AW performed enteroid experiments. IG provided critical material resources for this work. MC, EMH, YHT assembled figures. MC, EMH and JRS wrote the manuscript. EMH, YHT, AW, KDW, CC contributed methods. All authors edited, read and approved the manuscript.

## METHODS

### Isolating, establishing and maintaining human fetal enteroids

Fresh human fetal epithelium was isolated and maintained as previously described (Tsai et al., 2018). Once enteroids were established, healthy cystic enteroids were manually selected under a stereoscope and bulk-passaged through a 30G needle and embedded in Matrigel (Corning, 354234). For single-cell passaging, healthy cystic enteroids were manually selected under a stereoscope and dissociated with TrypLE Express (Gibco, 12605-010) at 37°C before filtering through 40μm cell strainers. Cells were then counted using a hemocytometer (ThermoFisher) and embedded in Matrigel.

### Media composition

Culture media consisted of 25% LWRN conditioned media generated as previously described (Miyoshi and Stappenbeck, 2013; Tsai et al., 2018) and 75% Human 2X basal media [Advanced DMEM/F12 (Gibco, 12634-028); Glutamax 4 mM (Gibco, 35050-061); HEPES 20 mM (Gibco, 15630-080); N2 Supplement (2X) (Gibco, 17502-048), B27 Supplement (2X) (17504-044), Penicillin-Streptomycin (2X) (Gibco, 15140-122), N-acetylcysteine (2 mM) (Sigma, A9165-25G), Nicotinamide (20 mM) (Sigma, N0636-061)]. This culture media was the base media for the eight culture conditions with varied concentrations of rhEGF (R&D, 236-EG) and rhNRG1 (R&D, 5898-NR-050) as follows: 100 ng/mL EGF with 0, 1, 10, and 100 ng/mL NRG1; 100 ng/mL NRG1 with 0, 1, and 10 ng/mL EGF; and culture media with neither EGF nor NRG1.

### Human subjects

Normal, de-identified human fetal intestinal tissue was obtained from the University of Washington Laboratory of Developmental Biology. All human tissue used in this work was de-identified and was conducted with approval from the University of Michigan IRB

### Single cell dissociation

To dissociate human fetal tissue to single cells, fetal duodenum was first dissected using forceps and a scalpel in a petri dish filled with ice-cold 1X HBSS (with Mg^2+^, Ca^2+^). Whole thickness intestine was cut into small pieces and transferred to a 15 mL conical tube with 1% BSA in HBSS. Dissociation enzymes and reagents from the Neural Tissue Dissociation Kit (Miltenyi, 130-092-628) were used, and all incubation steps were carried out in a refrigerated centrifuge pre-chilled to 10°C unless otherwise stated. All tubes and pipette tips used to handle cell suspensions were pre-washed with 1% BSA in 1X HBSS to prevent adhesion of cells to the plastic. Tissue was treated for 15 minutes at 10°C with Mix 1 and then incubated for 10 minute increments at 10°C with Mix 2 interrupted by agitation by pipetting with a P1000 pipette until fully dissociated. Cells were filtered through a 70µm filter coated with 1% BSA in 1X HBSS, spun down at 500g for 5 minutes at 10°C and resuspended in 500µl 1X HBSS (with Mg^2+^, Ca^2+^). 1 mL Red Blood Cell Lysis buffer was then added to the tube and the cell mixture was placed on a rocker for 15 minutes in the cold room (4°C). Cells were spun down (500g for 5 minutes at 10°C), and washed twice by suspension in 2 mL of HBSS + 1% BSA, followed by centrifugation. Cells were counted using a hemocytometer, then spun down and resuspended to reach a concentration of 1000 cells/μL and kept on ice. Single cell libraries were immediately prepared on the 10x Chromium at the University of Michigan Sequencing Core facility with a target of 5000 cells. The same protocol was used for single cell dissociation of healthy cystic enteroids manually collected under a stereoscope. A full, detailed protocol of tissue dissociation for single cell RNA sequencing can be found at www.jasonspencelab.com/protocols.

### Single cell library preparation

All single-cell RNA-seq sample libraries were prepared with the 10x Chromium Controller using either the v2 or v3 chemistry. Sequencing was performed on an Illumina HiSeq 4000 or NovaSeq with targeted depth of 100,000 reads per cell. Initial cell demultiplexing and gene quantification were performed with the default 10x Cellranger pipeline using the pre-prepared hg19 reference.

### Primary tissue collection, fixation and paraffin processing

Human fetal intestine tissue samples were collected as ~0.5 cm fragments and fixed for 24 hours at room temperature in 10% Neutral Buffered Formalin (NBF), and washed with UltraPure Distilled Water (Invitrogen, 10977-015) for 3 changes for a total of 2 hours. Tissue was dehydrated by an alcohol series diluted in UltraPure Distilled Water (Invitrogen, 10977-015).

Tissue was incubated for 60 minutes each solution: 25% Methanol, 50% Methanol, 75% Methanol, 100% Methanol. Tissue was stored long-term in 100% Methanol at 4°C. Prior to paraffin embedding, tissue was equilibrated in 100% Ethanol for an hour, and then 70% Ethanol. Tissue was processed into paraffin blocks in an automated tissue processor (Leica ASP300) with 1 hour changes overnight.

### Multiplex Fluorescent *In Situ* Hybridization (FISH)

Paraffin blocks were sectioned to generate 5 μm-thick sections within a week prior to performing *in situ* hybridization. All materials, including the microtome and blade, were sprayed with RNase-away solution prior to use. Slides were baked for 1 hour in a 60°C dry oven the night before, and stored overnight at room temperature in a slide box with a silicone desiccator packet, and with seams sealed using parafilm. The in situ hybridization protocol was performed according to the manufacturer’s instructions (ACD; RNAscope multiplex fluorescent manual protocol, 323100-USM) under standard antigen retrieval conditions and 30 minute protease treatment. Immediately following the HRP blocking for the C2 channel of the FISH, slides were washed three times for 5 minutes in PBS, then transferred to blocking solution (5% Normal Donkey Serum in PBS with 0.1%Tween-20) for 1 hour at room temperature. Slides were then incubated in primary antibodies overnight at 4°C in a humidity chamber. The following day, excess primary antibodies were rinsed off through a series of PBS washes. Secondary antibodies and DAPI (1 μg/ml) were added and slides were incubated at room temperature for 1 hour. Excess secondary antibodies were rinsed off through a series of PBS washes, and slides were mounted in ProLong Gold (TermoFisher, P36930). A list of antibodies and concentrations can be found in the Key Resources Table. All imaging was done using a NIKON A1 confocal and images were assembled using Photoshop CC. Z-stack series were captured and compiled into maximum intensity projections using NIS-Elements (Nikon). Imaging parameters were kept consistent for all images within the same experiment and any post-imaging manipulations were performed equally on all images from a single experiment.

### Single-cell in silico analysis

All in silico analyses downstream of gene quantification was done using Scanpy with the 10x Cellranger derived gene by cell matrices (Wolf et al., 2018). All samples were filtered to remove cells with less than 1000 or greater than 9000 genes, and less than 3500 or greater than 25000 counts per cell. Raw read counts per gene were scaled and log normalized prior to analysis.

Fetal tissue samples were batch corrected using BBKNN prior to dimensional reduction by principal component calculation and UMAP (McInnes et al., 2018; Polański et al., 2019). Genes were not included in the analysis if they were not sufficiently statistically invariable between cells. Clusters of cells within the combined full time-course of data were calculated using the Louvain algorithm within Scanpy with a resolution of 0.6. Cell type scoring was done with the native Scanpy score_genes() scoring function, as previously reported. Fetal tissue cell type scoring was conducted based on gene lists of established markers for each cell type and newly defined markers of submucosal and subepithelial cells (see supplemental data for gene lists)(Satija et al., 2015). Cell type scoring for *in vitro* grown enteroids was done based on gene lists derived from human homologs of cell type specific gene lists from Haber et al., 2017 (Haber et al., 2017) (see supplemental data for enteroids cell type gene lists).

## DATA AND CODE AVAILABILITY

All code used for single cell analysis and data presentation is available via Github at: (https://github.com/jason-spence-lab/fetal_intestine). Submission of raw sequencing data is in process, for inquiries contact authors.

