## Supplementary figures and images for "*In vitro* and *in vivo* development of the human intestinal niche at single cell resolution"

### Supplemental Figure 1

Supplemental Figure 1

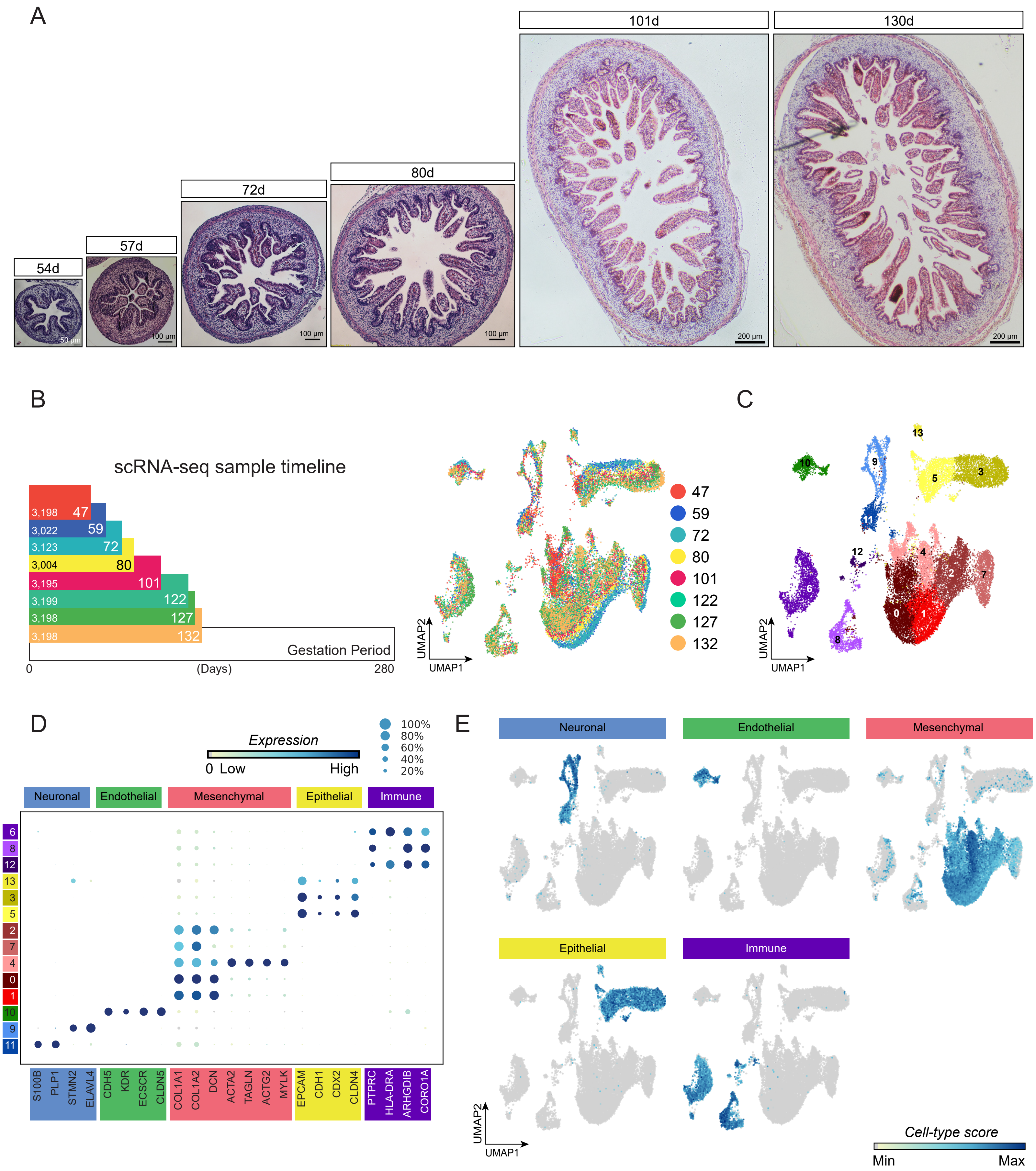

### Supplemental Figure 2

Supplemental Figure 2

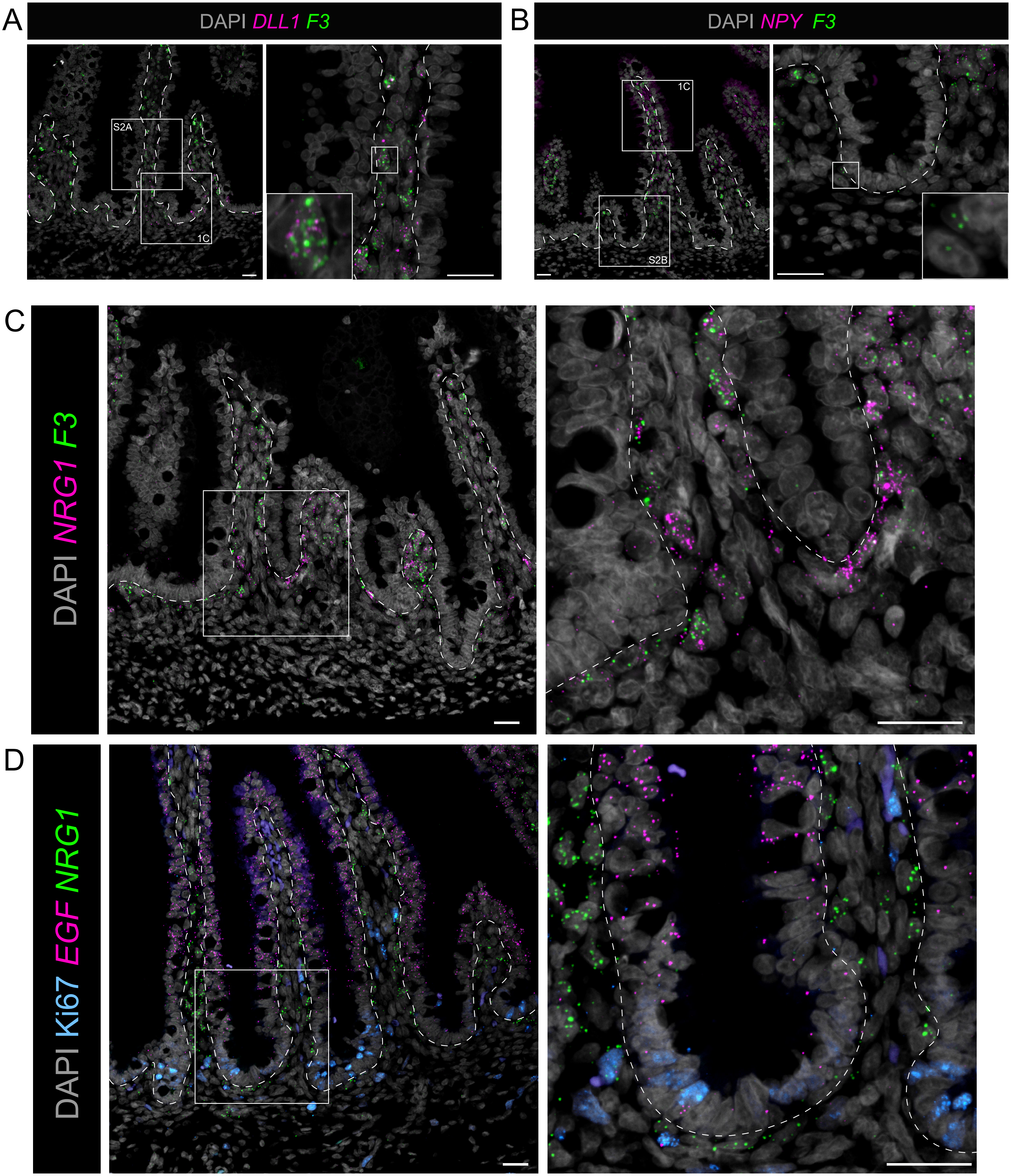

### Supplemental Figure 3

#### Supplemental Figure 3

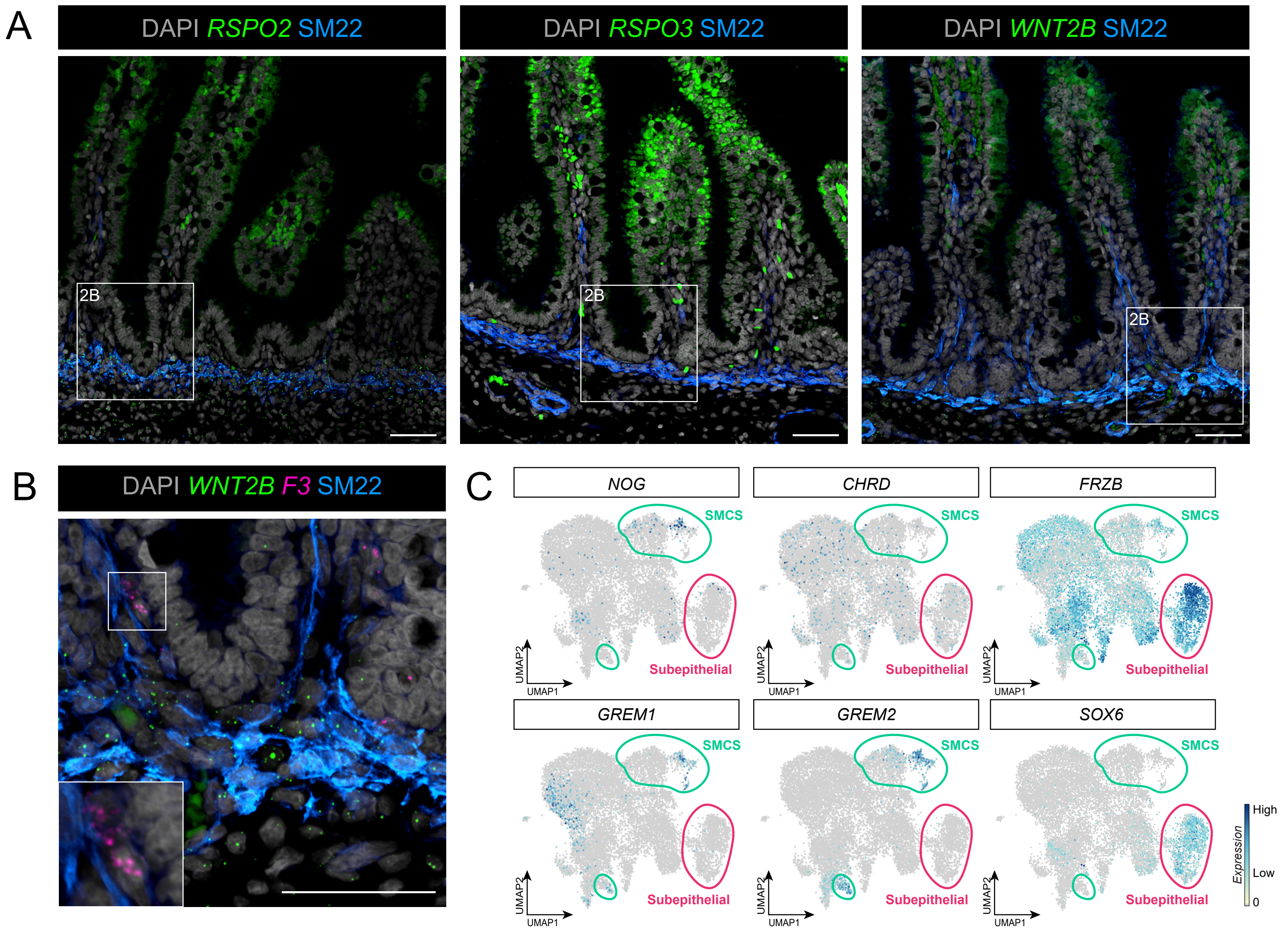
